# Yeast9: A Consensus Yeast Metabolic Model Enables Quantitative Analysis of Cellular Metabolism By Incorporating Big Data

**DOI:** 10.1101/2023.12.03.569754

**Authors:** Chengyu Zhang, Benjamín J. Sánchez, Feiran Li, Cheng Wei Quan Eiden, William T. Scott, Ulf W. Liebal, Lars M. Blank, Hendrik G. Mengers, Mihail Anton, Albert Tafur Rangel, Sebastián N. Mendoza, Lixin Zhang, Jens Nielsen, Hongzhong Lu, Eduard J. Kerkhoven

**Affiliations:** State Key Laboratory of Microbial Metabolism, School of Life Sciences and Biotechnology, Shanghai Jiao Tong University, Shanghai 200240, China; State Key Laboratory of Bioreactor Engineering, and School of Biotechnology, East China University of Science and Technology (ECUST), Shanghai, 200237, China; The Novo Nordisk Foundation Center for Biosustainability, Technical University of Denmark, DK-2800 Kgs Lyngby, Denmark; Department of Biotechnology and Biomedicine, Technical University of Denmark, DK-2800 Kgs Lyngby, Denmark; Institute of Biopharmaceutical and Health Engineering, Tsinghua Shenzhen International Graduate School, Tsinghua University, Shenzhen 518055, China; School of Chemistry, Chemical Engineering and Biotechnology, Nanyang Technological University, 62 Nanyang Drive, Singapore 637459; UNLOCK, Wageningen University & Research, Wageningen, The Netherlands; Laboratory of Systems and Synthetic Biology, Wageningen University & Research, Wageningen, The Netherlands; Institute of Applied Microbiology - iAMB, Aachen Biology and Biotechnology - ABBt, RWTH Aachen University, 52074 Aachen, Germany; Department of Life Sciences, National Bioinformatics Infrastructure Sweden, Science for Life Laboratory, Chalmers University of Technology, Gothenburg SE412 58, Sweden; Department of Life Sciences, Chalmers University of Technology, Gothenburg SE412 96, Sweden; Center for Mathematical Modeling, University of Chile, Santiago, Chile; Systems Biology Lab, Vrije Universiteit Amsterdam, Amsterdam, the Netherlands; BioInnovation Institute, Ole Maaløes Vej 3, DK2200 Copenhagen N, Denmark; Department of Life Sciences, SciLifeLab, Chalmers University of Technology, Gothenburg SE412 96, Sweden

**Keywords:** *Saccharomyces cerevisiae*, Genome-scale metabolic models, Machine learning, Multi-omics integration

## Abstract

Genome-scale metabolic models (GEMs) can facilitate metabolism-focused multi-omics integrative analysis. Since Yeast8, the yeast-GEM of *Saccharomyces cerevisiae*, published in 2019, has been continuously updated by the community. This have increased the quality and scope of this model, culminating now in Yeast9. To evaluate its predictive performance, we generated 163 condition-specific GEMs constrained by single-cell transcriptomics from osmotic pressure or normal conditions. Comparative flux analysis showed that yeast adapting to high osmotic pressure benefits from upregulating fluxes through the central carbon metabolism. Furthermore, combining Yeast9 with proteomics revealed metabolic rewiring underlying its preference in nitrogen sources. Lastly, we created strain-specific GEMs (ssGEMs) constrained by transcriptomics for 1229 mutant strains. Well able to predict the strains’ growth rates, fluxomics from those large-scale ssGEMs outperformed transcriptomics in predicting functional categories for all studied genes in machine-learning models. Based on those findings we anticipate that Yeast9 will empower systems biology studies of yeast metabolism.

## Introduction

*Saccharomyces cerevisiae*, a widely used model organism in eukaryotic studies, was the first eukaryote whose genome was thoroughly sequenced and annotated ^1^. *S. cerevisiae* has long been used to study genetic interactions ^2,3^, build cell factories for the production of high-value-added compounds ^4–6^, and comprehend eukaryotic metabolism due to its clear genetic background, abundant gene annotation and genetic tractability ^7–9^. Having amassed extensive knowledge of the metabolism and physiology of yeast such as *S. cerevisiae*, the genome-scale metabolic models (GEMs) of this organism have undergone 20 years of iterative refinement and enhancement including Yeast8^10^, Yeast7^11^, Yeast6^12^, iND750^13^, iLL672^14^ since the initial publication of the first-generation model, iFF708, in 2003 ^15^. The establishment and maturation of yeast-GEMs have laid a strong foundation for the emergence of a model ecosystem centered on *S. cerevisiae*, including ecYeast, etcYeast, and pcYeast, collectively enabling a variety of system and synthetic biology investigations concerning *S. cerevisiae* ^10,16^. For instance, leveraging the enforced objective flux (FSEOF) algorithm ^17^, Yeast8 and ecYeast8 has been used to obtain a 70-fold improvement in heme production ^18^. Kinetic models deprived from yeast-GEM possess the capability to discern species-specific details employing the kinetic parameterization procedure ^19–21^.

The ability by which yeast, like any other cellular biocatalyst, can overproduce desirable chemicals and secrete commercial proteins depends on cellular gene expression, which is determined by both the genotype and the environment. Diverse growth environments, encompassing variations in nutrition, temperature and stress, possess the capacity to strongly shape cellular metabolism, thereby exerting significant influence on the phenotypic outputs. Contrastingly, traditional GEMs are is chiefly based on the whole genome sequence and their functional gene annotations ^22^. As a result, it is challenging for GEMs themselves to reflect gene expression levels corresponding to dynamic environmental changes. With the accumulation of various omic datasets, there has been a growing interest in reconstruction of omics-constrained GEMs, especially in *Escherichia coli*, where the biological activity of metabolic networks is contextualized based on quantitative transcriptomics or proteomics ^23–28^. By contrast to ordinary GEM, those context-specific GEMs are more powerful in simulating and revealing metabolic changes under environmental and genetic perturbations. However, until now, few studies have been conducted to systematically evaluate the quality and prediction capabilities of large-scale context-specific yeast-GEMs. On the other hand, building and analyzing context-specific GEMs needs more accurate gene-protein-reaction relationships (GPR), protein compartment annotation, thermodynamics information, etc. However, yeast-GEMs are still deficient in the above aspects, making the systematic reconstruction and analysis of yeast context-specific GEMs laborious.

To fill the aforementioned gaps in Yeast8 and its derived condition-specific metabolic models, we released the latest version of the yeast-GEM (Yeast9) for the community by merging consistent model updates that have been made in the past four years. To display the unique value of Yeast9 in transforming big data into knowledge, through leveraging the large-scale omics and phenotype datasets, we systematically reconstructed and analyzed numerous omics-constrained GEMs derived from Yeast9, i.e., 163 condition-specific GEMs (csGEMs) at single-cell level to decipher the metabolic readaptation mechanism under high osmotic stress and 1,143 strain-specific GEMs (ssGEMs) to characterize the yeast metabolism under single gene deletion. These studies showcase that Yeast9 is well suited for conducting omics integrative analysis in order to uncover complex relations between genotype and phenotype. Moreover, yeast-GEMs can be used for predicting cellular responses to novel environmental conditions, as well as being computational platforms to pioneer the development of the industrial strains. Therefore, Yeast9 can serve as a computational toolbox for quantifying yeast physiology and guiding experimental works for the wider yeast community.

## Results

### Model improvements from the yeast community

Through a collective effort of iterative engagements by the yeast-GEM community, the consensus yeast-GEM was updated from Yeast8 to Yeast9. Following a similar pipeline as employed for Yeast8, every round of updates was diligently recorded and comprehensively documented through a version-controlled system (https://GitHub.com/SysBioChalmers/yeast-GEM) which provided transparency and reproducibility in the development of Yeast9. The coverage of the metabolic network was increased by a combination of targeted expansion by, e.g., including reactions related to volatile esters & polyphosphates, and by identification of missing gene-protein-reaction relations (GPRs) from reaction databases KEGG ^29^ and MetaCyc^30^. Combined with further curations described below, this yielded 30 new genes, 203 new reactions, and 140 new metabolites (Fig. 1).

**Figure 1.**
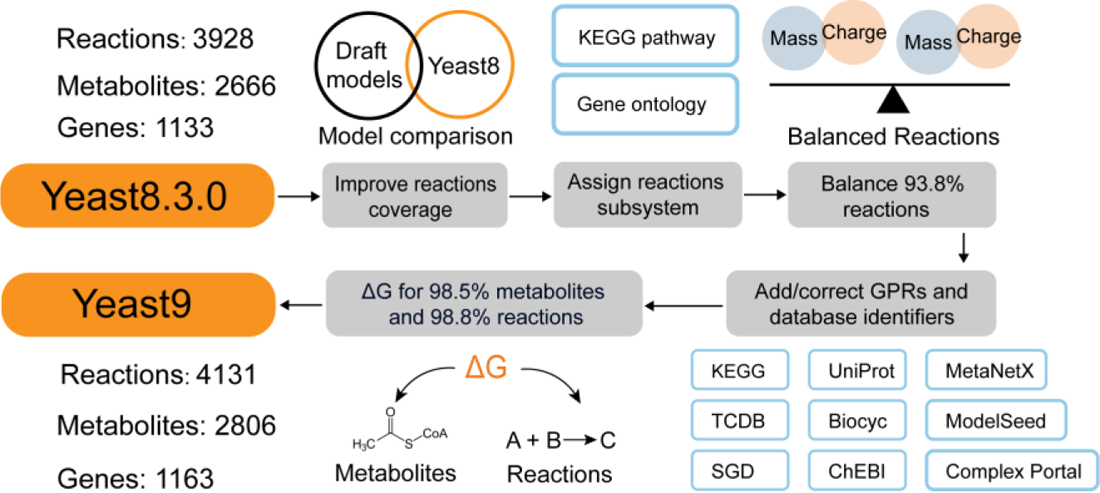
Major improvements in Yeast9 compared to Yeast8. The Yeast9 contains 1,163 genes, 2,806 metabolites, and 4,131 reactions. New reactions were identified by comparing Yeast8 with draft models constructed by RAVEN. ΔG°’ was added for almost all metabolites and reactions. Each reaction was linked with a single subsystem according to the pathway annotation from KEGG or SGD. Various GPRs were added or corrected by multi-rounds of manual comparison with databases. Nearly all reactions were curated to ensure mass and charge balances.

Numerous GPRs and metabolite annotations were curated by reviewing the corresponding annotation from NCBI ^31^, UniProt ^32^, KEGG, ChEBI ^33^, PubChem ^34^, MetaNetX ^35^, ModelSeed ^36^, BiGG ^37^, BioCyc ^30^ and Reactome ^38^. We systematically curated all annotations of transport reactions according to the detailed protein function annotations at SGD ^39^, TCDB ^40^, YeastCyc ^30^, KEGG, and UniProt (Fig. 1), lending confidence to simulations involving transporter usage. The subunit composition of 36 protein complexes was corrected based on SGD, Complex Portal^41^ and UniProt. To facilitate pathway analyses, each reaction was assigned to single explicit subsystems, according to the subsystem annotations (Supplementary File 1) from the KEGG, BioCyc and the GO ontology in SGD. The top 20 subsystems were summarized in Supplementary Figure 1.

When simulating flux distributions with the updated model, the feasibility of metabolic fluxes and their directionality, can be determined by thermodynamics analysis. We assigned ΔG°’ for 98.5% metabolites and 98.8% reactions according to evidence gathered from the yETFL model ^42^, dGPredictor ^43^ and ModelSEED database. Furthermore, we balanced most mass/charge unbalanced reactions in the model, thus increasing the percentage of balanced reactions to 93.8%.

### Systematic evaluation of Yeast9 in its prediction quality

Through the above improvements, Yeast9 contains 2806 metabolites, 1163 genes, and 4131 reactions. The newly added thermodynamic information makes it possible to explore the driving forces of mass transformation in metabolism at both single reaction and sub-pathway levels. Figure 2a shows that the ΔG°’ in the central carbon metabolism, in which ΔG°’ of EMP is −22.8 kJ/mol, TCA is −9.6 kJ/mol and PPP is −13.8 kJ/mol, under biochemical standard conditions (pH = 7, ionic strength = 0.25 in cytoplasm and mitochondrion). Inspecting individual reactions demonstrates that some reactions (i.e., within the EMP pathway) with positive ΔG°’ are able to proceed in the forward direction regardless. Potential disparities between biochemical standard and true intracellular conditions would have some effect on the ΔG of reactions. Regardless, overcoming a large positive ΔG°’ is likely indicative of a ratio of reactants and products that is for away from the equilibrium constant of that reaction. This and the proximity of strongly negative ΔG°’ reactions yields overall negative ΔG°’ at the sub-pathway level, enabling enzymatic reactions even with positive ΔG°’.

**Figure 2.**
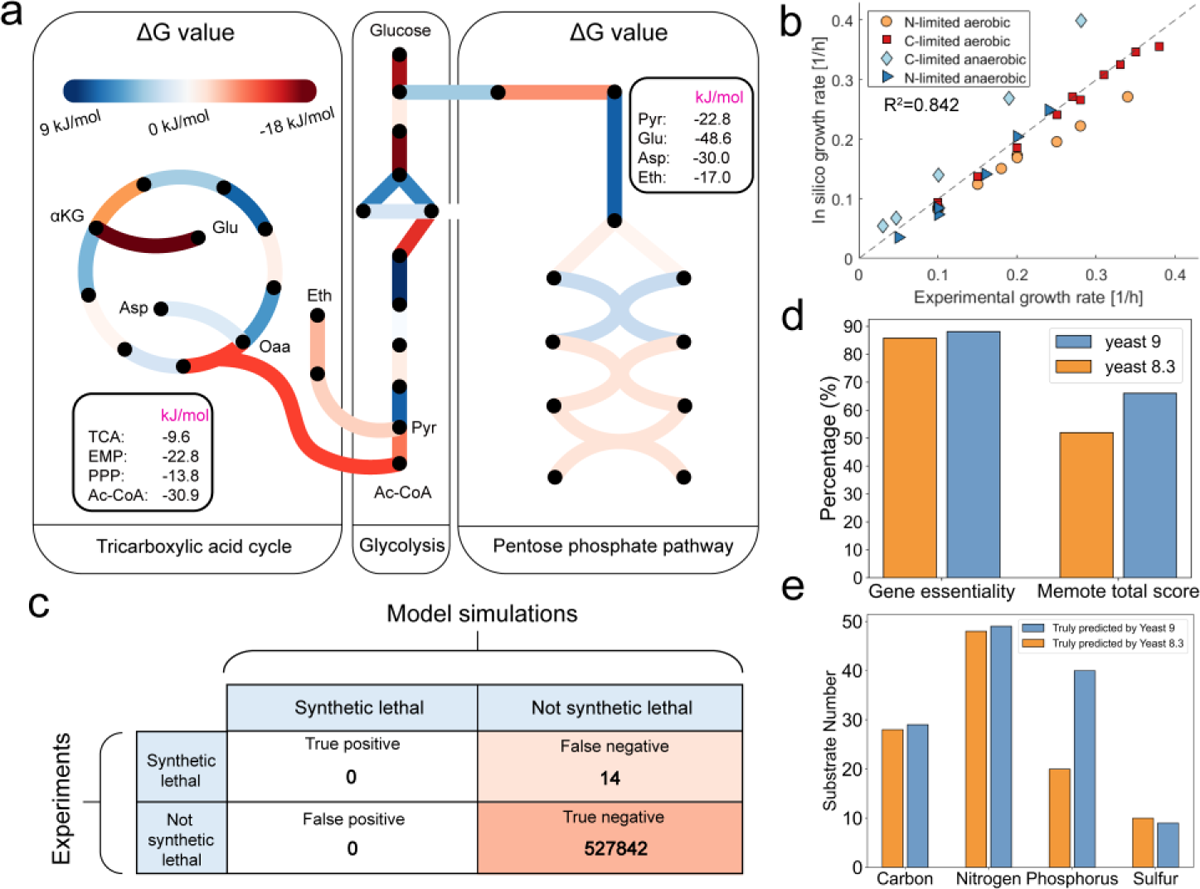
Systematic evaluation of prediction capability by Yeast9. (a) The profile of ΔG°’ in TCA, EMP and PPP. The color denotes ΔG°’ value. The red line means that the reactions are favorable thermodynamically; the blue line indicates that the reactions are unfavorable thermodynamically. The number within the bold rounded rectangle represents the total ΔG°’ of TCA, EMP, PPP and reaction pathways in synthesizing acetyl-CoA, pyruvate, glutamine, aspartate and ethanol from glucose. (b) Growth simulation under aerobic and anaerobic conditions. (c) The Yeast9 could predict the consequences of synthetic lethal of two gene combinations, with the accuracy almost at 100%. (d) Comparison in predicted gene essentiality and Memote score between Yeast8 and Yeast9. (e) Carbon, nitrogen, phosphorus and sulfur source usages comparison between Yeast8 and Yeast9.

Yeast9 has an improved performance in characterizing cell growth from a wide range of conditions compared to Yeast8. Predicted growth from Yeast9 correlated well with experimental data under both aerobic and anaerobic conditions with R^2^ equals to 0.842 (Fig. 2b). With 14% improvement in MEMOTE score^44^, the predictions of gene essentiality and substrate usage by Yeast9 were moderately improved compared with Yeast8 (Fig. 2d, 2e). Synthetic lethality can be predicted with near-perfect accuracy (Fig. 2c), whereas a few cases of false negative predictions still exist (Supplementary File 2). For example, the genes YNL192W and YLR342W code for chitin and glucan synthases respectively, both indispensable for a functional cell wall. Double-deletion of those genes are lethal in the laboratory ^3^, however, the relevant reactions in Yeast9 are annotated with redundant isozymes that prevent *in silico* lethality. *In vivo*, however, the expression of the isozymes are transcriptionally regulated ^45,46^, which is an aspect that is not considered in GEMs, and likely the explanation for the false negative prediction.

### Environmental adaptation mechanism revealed by single-cell omics-constrained GEMs derived from Yeast9

While yeast-GEM represents the theoretical metabolic network of *S. cerevisiae* based on its genome annotation, not all enzymes might be constitutively expressed. To examine this, we collected 163 single-cell transcriptomes of *S. cerevisiae* ^47^ including 80 transcriptomes measured under high osmotic stress and 83 transcriptomes measured under reference conditions. We used GIMME ^24^ to construct 163 single-cell omics-constrained GEMs (scGEMs) (Fig. 3a). The generated scGEMs have reaction numbers ranging from 2,278 to 3,858, with the numbers of metabolite ranging from 2,114 to 2,708.

**Figure 3.**
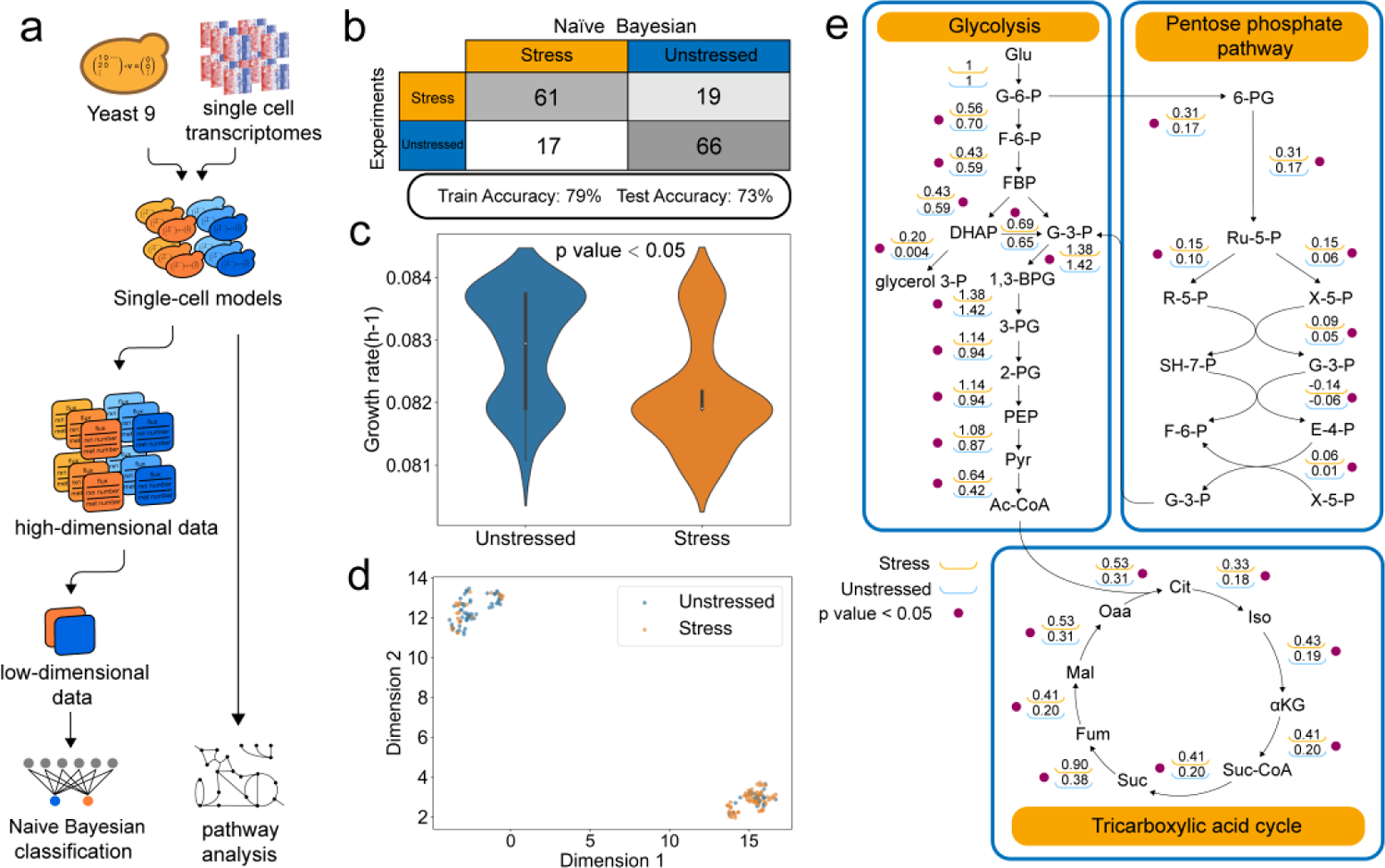
Construction and application of scGEMs derived from Yeast9. (a) Construction and analytic workflows. We integrated single-cell transcriptomics measured under osmotic stress conditions and normal conditions into yeast-GEM by GIMME, resulting in 163 scGEMs. (b) The Naïve Bayes classifier has 73% accuracy in classifying the single cell from high osmotic stress and normal conditions. (c) Predicted growth rates of single cells from osmotic stress conditions and normal conditions. (d) The UMAP is used to reduce the high-dimensional data into reduced data, with dimension 1 plotted against dimension 2. (e) Comparison of mean flux values for central carbon metabolism as predicted by condition-specific scGEMs.

We gathered the metabolite number, reaction number, and projected flux of each scGEM to categorize *S. cerevisiae* cells sampled from stressed and unstressed conditions (Fig. 3a). Uniform manifold approximation and projection (UMAP) was executed to extract features before machine learning ^48^. The Naïve Bayes classification was trained using the processed data. The Naïve Bayes classifier demonstrated 73% accuracy in differentiating single cell sampled from the osmotic stress and unstressed conditions (Fig. 3b). This implies that the metabolic networks of individual cells were sufficiently different to enable categorizing them by which environment they resided. To further analyze the mechanism by which *S. cerevisiae* responds to osmotic stress conditions, single-cell specific growth rates (Fig. 3c) and central carbon metabolism fluxes (Fig. 3e) were calculated by simulating the scGEMs. While generally *S. cerevisiae* grew slower under osmotic stress (*P* value < 0.05), a subset of stressed cells grew similarly to those in unstressed cells and vice versa (Fig. 3c). This shows a possible existence of a resistant phenotype or alternative stress response pathways that might be worth exploring. Cluster analysis on the simulated fluxes further illustrated high similarity within the same condition, while retaining heterogeneity (Fig. 3d, Supplementary Figure 2). In terms of evolution, heterogeneity at the single-cell level may assist *S. cerevisiae* populations to efficiently adapt to new environments.

In a more detailed analysis, we examined the flux distributions from stressed and unstressed cells, particularly focusing on the tricarboxylic acid cycle (TCA), pentose phosphate pathway (PPP), and glycolysis (Fig. 3e). About 46% of the active fluxes are divergent (*P* value < 0.05) between unstressed cells and stressed cells (Supplementary Figure 3), in central carbon metabolism all reactions were significantly distinct (*P* value < 0.05). An increased flux through PPP, TCA, and lower-glycolytic fluxes was found in stressed yeast cells, which is consistent with proteomics data ^49^, which show that the overall protein expression of these three pathways are changed in the same direction as the flux. This signifies that *S. cerevisiae* strengthens it central carbon metabolism to generate more ATP, provide NADPH redox potential, and possibly synthesize more precursors in response to high osmotic stress. Furthermore, the stressed scGEMs had an increased flux from dihydroxyacetone phosphate (DHAP) via glycerol 3-phosphate to glycerol, an osmoprotectants whose biosynthesis during osmostress is well documented ^50,51^.

### Multi-omics analysis based on Yeast9 elucidates the metabolic rewiring under nitrogen limitation

Another environmental parameter affecting metabolism is nutrient availability, and as has been reported, *S. cerevisiae* preferentially assimilates ammonium or certain amino acids, specifically glutamine and glutamate ^52,53^. To check whether Yeast9 could quantitatively classify the preferred nitrogen source utilized by *S. cerevisiae*, preference scores for the nitrogen sources ammonium, glutamate, isoleucine and phenylalanine were calculated using Yeast9 (Fig. 4a). Here, the nitrogen preference score is defined as the absolute value of the slope between nitrogen and glucose uptake rates, where the uptake of a non-preferred nitrogen source will result in a more significant increase in glucose uptake when compared to a preferred nitrogen source. The order of preferred nitrogen sources (Fig. 4b) based on Yeast9 predictions were consistent with previous studies ^53,54^.

**Figure 4.**
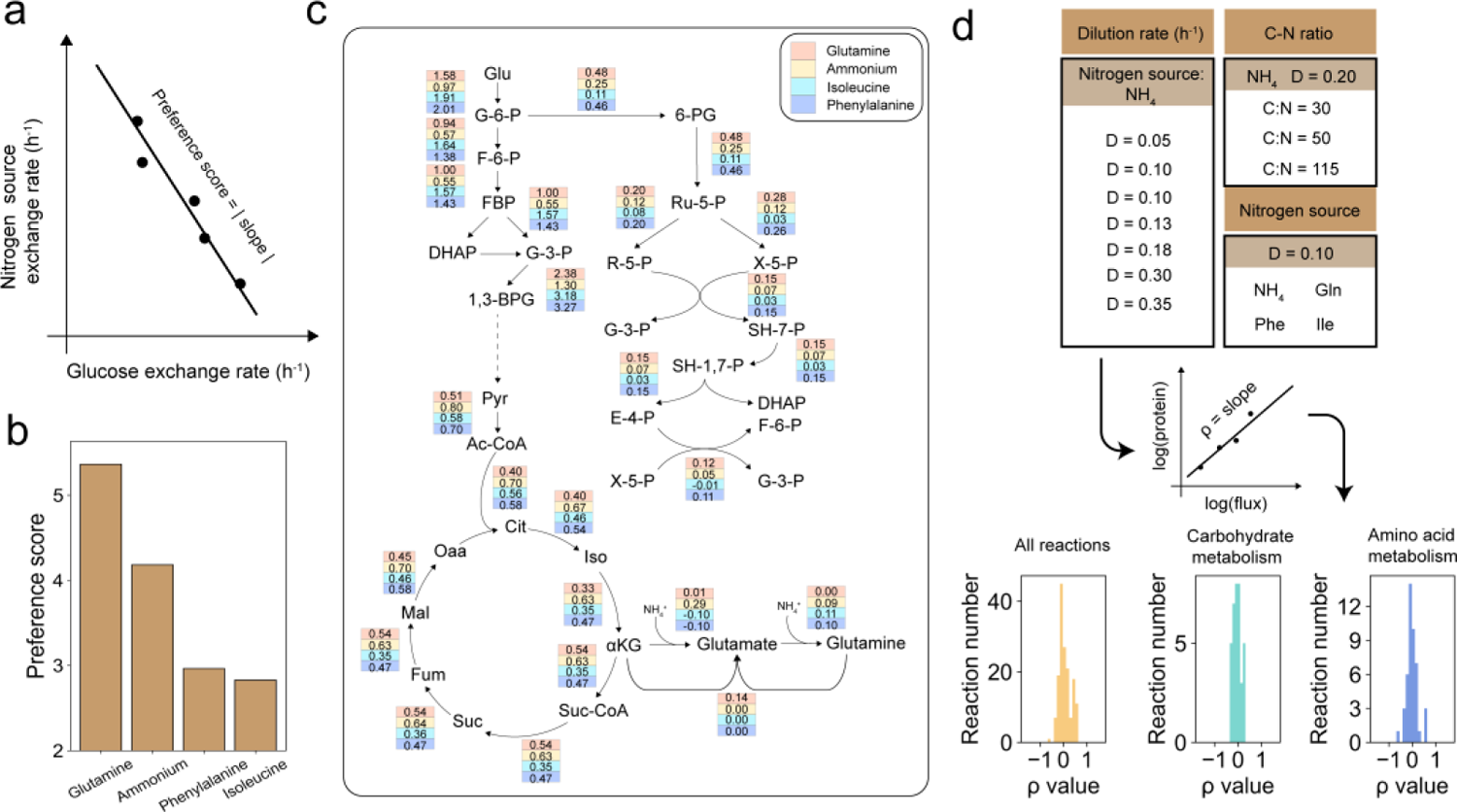
Illustration of yeast metabolism rewiring under nitrogen-limitation based on Yeast9 and multi-omic integrative analysis. (a) A graphical representation of the preference score. The preference score denotes the degree to which the change in glucose uptake rate compensates for the decrease in nitrogen source absorption rate. (b) The preference scores of glutamine, ammonium, isoleucine, and phenylalanine. (c) pFBA was used to get the flux distributions of 4 nitrogen sources. The differences between the 4 flux distributions were analyzed by Kruskal– Wallis test. For all reactions in (c), *P* values < 0.05. (d) 14 phenotype-constrained models and the calculation of protein regulation coefficient (ρ). The correlation analysis between protein abundances and fluxes was conducted at both holistic and sub-pathway levels.

To further investigate how nitrogen sources tune yeast metabolism, integrative analysis with Yeast9 was carried out by leveraging reported multi-layer omics datasets ^55,56^. As the first step, we calculated flux distributions by constraining Yeast9 with measured exchange fluxes and total protein concentrations. As shown in Fig. 4c, nitrogen sources largely reshape cellular flux distribution. Next, we analyzed consistent tendencies between reaction fluxes and the related protein abundances, comparing the unpreferred nitrogen sources (isoleucine and phenylalanine) with the favored one (i.e., ammonia). Regarding the genes covered by in Yeast9, 13.6% (159 for isoleucine) and 23.2% genes (270 for phenylalanine) show consistent fluctuations in both flux and protein level (Supplementary File 3), which suggests that these reactions are transcriptionally regulated. There are 145 common genes selected from the above two groups of genes, which were enriched in amino acid biosynthesis (*P* value < 0.001) by GO term enrichment analysis (Supplementary Fig. 4).

To more precisely determine the covariance between fluxomics and proteomics across diverse environments, we further conducted regulatory analysis by utilizing 14 phenotype-constrained models (Fig. 4d) and the associated proteomic datasets, as described previously ^57^. The contribution of gene expression to flux can be calculated as the slope between log(flux value) and log(protein abundance) (Fig. 4d), which was defined as the so-called protein regulation coefficient (ρ) ^58,59^. Here, ρ ≈ 1 signifies that changes in simulated fluxes can largely be explained by protein concentration changes. Most reactions from carbohydrate metabolism and amino acid metabolism exhibited weak protein regulation coefficients (Fig. 4d), with only 11 reactions revealing high coefficients (ρ > 0.5, Supplementary file 4). Overall, the weak correlation between proteomics and fluxomics illustrates that gene expressional changes alone do not fully reflect flux changes. As proteome changes are not necessarily reflected in flux changes, multi-omics analyses are rendered more valuable to reflect metabolic adaption mechanisms.

### Growth profiles and gene function can be predicted by Yeast9 constrained with large-scale transcriptomic

When not lethal, gene knockouts may still alter gene expression levels, flux distributions and thereby phenotypes of an organism. Meanwhile, the knockout of genes that have similar functions may also yield similar changes in flux distributions. Therefore, it is possible to explore gene functions by analysing model-predicted fluxes and gene expression levels upon gene knockouts. To this purpose, we collected two transcriptomic datasets containing 1143 single knockout *S. cerevisiae* strains ^60,61^ and 86 single or double knockout strains to build 1229 ssGEMs using the early described method ^62^, where reaction bounds are changed according to gene expression levels (Fig. 5a). Flux balance analysis (FBA) with growth maximization as the objective function showed a good correlation between the predicted and measured growth rate (Pearson correlation coefficient = 0.66 and P < 0.05, Fig. 5b). Those inconsistent datapoints may be caused by the complex cellular regulation which were currently not captured by Yeast9.

**Figure 5.**
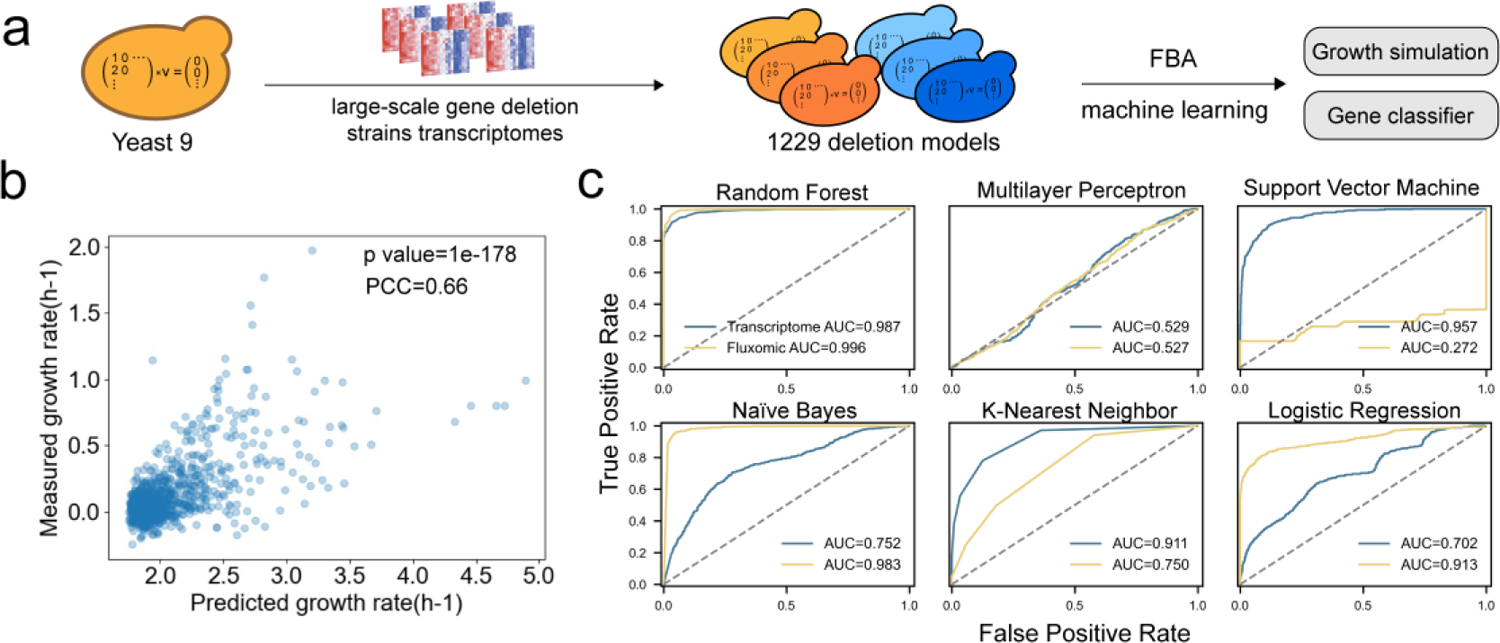
Large-scale transcriptomics-constrained GEMs built from Yeast9 could characterize the growth profiles of gene knock-out strains and classify gene functions. (a) The Yeast9 is constrained by transcriptomics to generate ssGEMs for gene knock-out strains, which are sequentially were used to predict the growth rate and train machine learning models in gene function classification. (b) Significant correlations existed between the relative doubling time (calculated based on measured value) and the corresponding simulated growth rate based on ssGEMs. (c) Fluxes and transcriptomics were used to train six machine learning algorithms to classify the genes into five different functional categories. The ROC and AUC of six classifiers using flux data (yellow line) or transcriptomic data (blue line) were computed by sklearn (https://scikit-learn.org/).

Previous studies have annotated 16 functional bioprocess categories (i.e., cell cycle regulation including YOR058C and YDR022C) encompassing the above-mentioned 1143 knocked-out genes ^62–66^ (Supplementary File 5). Using fluxomics or transcriptomics data as training dataset, six machine learning classifiers were separately trained to predict the function categories, to which those genes belong (Fig. 5c). The receiver operating characteristic curve (ROC) and area under the curve (AUC) show that, except for multilayer perceptron (MLP), the classifiers performed better when utilizing fluxomic as the only input compared to transcriptomics. We found utilizing both fluxomics and transcripomics as input, the prediction accuracy could not be improved further (Supplementary File 5). Among those 6 machine learning algorithms, the Naïve Bayes classifier with fluxomics as input showed the best performance and achieved relatively high accuracy for the training dataset (0.933) but also the testing dataset (0.874), together with high a recall rate (0.883) and F score (0.873). In conclusion, fluxes from the context-specific Yeast9 models provide sufficient information to classify gene function related to various cellular metabolic activities.

**Table 1.**
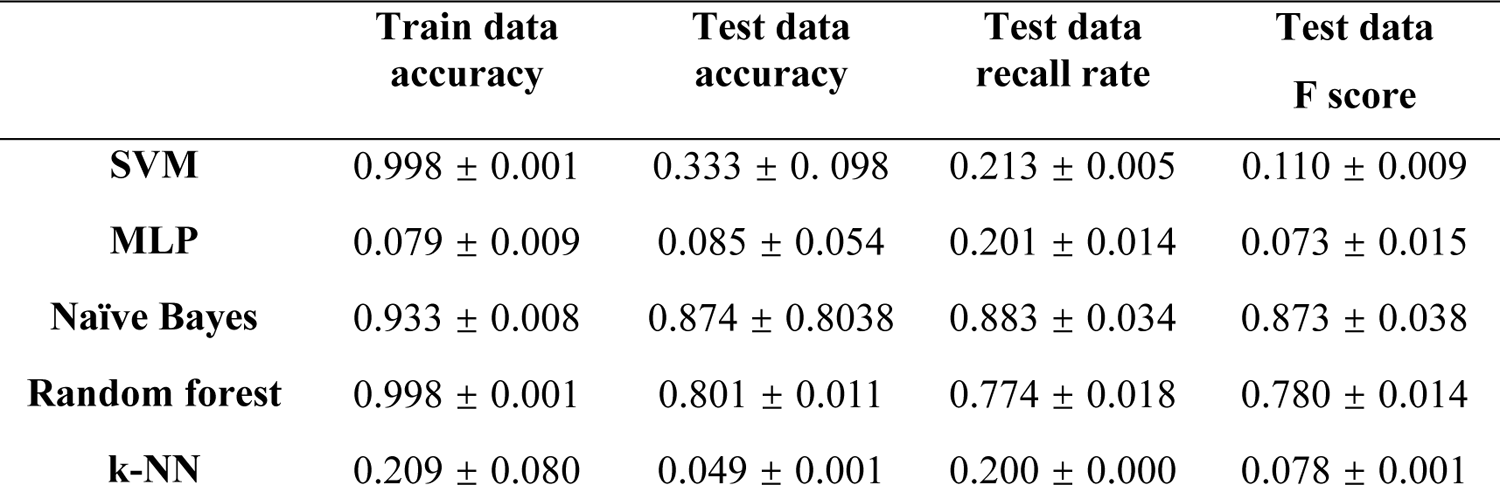
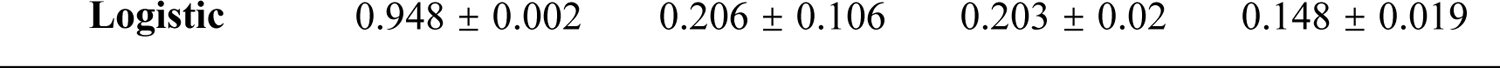
Summary of accuracy, recall rate, and F score of classifiers when using fluxomics as training datasets. Different machine learning models were trained to predict the classification of genes into the main gene functional categories with more than 70 genes. The fluxomics was processed by kPCA, which contains 500 components.

## Discussion

Through a concerted effort by members of the research community, we have updated yeast-GEM from Yeast8 to Yeast9, assisted by a version control system. The gradual improvements of yeast-GEMs covered a wide range of curations, including but not limited to the assignment of new gene functions; adjustment of incorrect gene assignments; modification of reaction directionality; and inclusion of annotations of proteins, metabolites and reactions. The abovementioned progress has filled gaps in current yeast-GEMs and enhanced their performance in comprehensively characterizing the yeast cellular metabolism. However, some limitations still exist for Yeast9. For example, it still lacks high-resolution details in representing yeast metabolic activities at the organelle level and some reactions for lipid synthesis are not standard. Thus, further efforts from the yeast community still need to be taken to refine the quality of Yeast9.

To solely rely on conventional GEM simulations for *in silico* studies of metabolism neglect the effect of mRNA and protein levels, leading to deviations between model simulations and real metabolic activities. We therefore evaluated Yeast9 in a number of integrative multi-omics analyses that quantified yeast physiology and metabolism. As the first example, 163 scGEMs could be classified according to their exposure to osmotic stress and showed subpopulations when considering their metabolic networks. Secondly, Yeast9 was able to enumerate which nitrogen sources (i.e., glutamine and ammonium) are most preferred by yeast. In additional simulations with Yeast9, different nitrogen sources could drastically alter the fluxes through central carbon metabolism. When considering integrative multi-layer omics analyses with Yeast9, we observed that a low consistency between changes in fluxomics and proteomics. Such a lower correlation has also been reported in other microorganisms, e.g., *E. coli* and *Bacillus subtilis* ^59,67–69^. Thereby, multi-omics analyses and GEM simulations are highly complementary when investigating metabolic regulation.

Overall, Yeast9 is the most comprehensive and state-of-the-art *S. cerevisiae* GEM as well as a valuable knowledge database on its metabolic network. Through continuing iterative updates, the quality of yeast-GEMs has further improved and this as the model can therefore function as an excellent template for generating high-quality GEMs for other non-conventional yeast species, such as *Pichia pastoris*, *Ogataea polymorpha* and Methylotrophic yeasts ^70–73^. Those models together can enable the exploration of evolutionary mechanisms that underly diverse metabolic activities and traits across yeast species. Ultimately, we are confident that the latest version of yeast-GEMs - Yeast9 and a flourishing model ecosystem, which still needs research community to contribute to further improvements, fostering an open-source and collaborative environment, around it will accelerate the developments in systems and synthetic biology studies of yeast in the coming years.

## Methods and materials

### Model curation

Aimed at adding new reactions and metabolites, two draft models were reconstructed using RAVEN Toolbox 2.0 ^74^. The two draft models were built based on KEGG and MetaCyc separately ^29,30^. Then, new reactions and metabolites were extracted by semi-automatically comparing the Yeast8 with those two draft models, after which the detailed annotation of genes, metabolites and reactions from MetaCyc, Yeastcyc, UniProt, SGD, and KEGG were utilized to guarantee that the new reactions and metabolites were reasonable and of high quality. Standard procedures for metabolites and reactions annotation used in this work were consistent with Yeast8 ^10^. In all, 203 new reactions and 140 new metabolites were added into Yeast9.

Reaction subsystems were systematically obtained from the KEGG pathway and integrated into the Yeast9. If KEGG does not have the subsystem annotation of the reactions, the Gene Ontology of the corresponding genes in SGD or basic biochemistry knowledge was applied. If no detailed information is found, the reactions are classified into the “other” subsystem.

Based on protein location data, the reactions in the lipid particle and peroxisome were curated. Taking the peroxisome as an example, if a protein is located in the peroxisome, but the peroxisome compartment in the current Yeast8 does not cover the related reaction, then the reaction is added into the peroxisome. On the contrary, if a protein is not located in the peroxisome, but the peroxisome compartment in the current Yeast8 has the corresponding reaction, then the reaction is removed from the peroxisome compartment. In terms of protein complexes’ location, the compartment annotation of all the subunits should be considered.

The Gibbs free energy change (ΔG°’) was added into yeast-GEM. The ΔG°’ for each reaction and metabolite is the same with yETFL model ^42^. The ΔG°’ of the reactions that were not contained in yETFL model is computed by dGPredictor ^43^. The ΔG°’ of the metabolites that were not in yETFL model is from Modelseed database.

The complex annotations were comprehensively refined, mainly based on ComplexPortal, and information from SGD and UniProt were also used. Additionally, the transporter annotations were curated using various databases, including TCDB, SGD, UniProt, and KEGG.

### Synthetic lethal simulation

Gene interactions data ^2,3,75–80^ were collected and the function “double_gene_deletion” in COBRA Toolbox was used to estimate synthetic lethal. Only the genes contained in the yeast-GEM were considered.

### scGEMs construction and simulation

Gasch et al previously described a series of single-cell transcriptomes of *S. cerevisiae*, in which 80 transcriptomes were measured under the osmotic stress condition and 83 transcriptomes under the unstressed condition ^47^. Constrained by the single-cell transcriptome dataset, the 163 single-cell-specific models were generated using the GIMME algorithm by COBRA Toolbox ^24^ in MATLAB based on Yeast9.

To analyse the metabolic difference between the osmotic stress condition and unstressed condition, parsimonious flux balance analysis (pFBA) ^81^ was used to maximise growth rate under the minimal medium condition. The resulting 163 flux distributions, together with the reaction numbers and metabolite numbers, formed 163 datasets. Secondly, dimensionality reduction was conducted by UMAP (components=3) for the Naïve Bayes classification. Then Naïve Bayes was performed by sklearn in Python, in which the dataset was randomly split into training (80%) and test (20%) datasets. The reduced data is also used for cluster analysis.

### Condition-specific GEMs construction and simulation under nitrogen limitation

Yeast9 was constrained by growth rate and exchange rate except for nitrogen exchange rate under nitrogen limitation condition^54,56^. The carbohydrate, protein, and RNA ratios in the biomass composition were scaled according to the measured carbohydrate, protein, and RNA abundance in the paper. As a result, 14 phenotype-constrained models were generated.

The preference score of yeast for different nitrogen sources was computed as follows:

- Step 1: Allow for uptake of one nitrogen source at the time, with a fixed growth rate of 0.1 h^-1^.
- Step 2: Determine the minimum nitrogen uptake by FBA while setting the relevant nitrogen source exchange reaction as objective function.
- Step 3: Step-wise (in 5 steps) increase the nitrogen uptake from 100% to 150% of the value determined in step 2, and determine the minimum glucose uptake by FBA while setting the glucose exchange reaction as objective function.
- Step 4: Determine the slope of glucose uptake versus nitrogen source uptake, and take its absolute value to represent the preference score.

To get the flux distribution under 4 nitrogen sources, Yeast9 was firstly constrained by the measured fluxes (multiplied by 0.8 considering the possible experimental error) for each related exchange reaction and the measured growth rate, with minimization of nitrogen source utilization as the objective function. The compositions of biomass in the model were refined by the measured ratios of protein, RNA under each condition. Then, the minimal nitrogen source uptake rate was calculated using FBA. Subsequently, the uptake rate of nitrogen source in models was fixed, to smaller than 1.2 times of the calculated minimal nitrogen source uptake rate. Afterwards, all fluxes were recalculated by pFBA, which was used in Supplemental Fig.5.

For genes selection in Supplemental Fig.5, the genes with higher protein expression level and corresponding flux than their counterpart in the model using NH4 as the only nitrogen source were selected. For genes related to more than one reaction, at least one related reaction flux larger than that in the NH4 model meets screening requirements. After that, The GO enrichment analysis (https://david.ncifcrf.gov/) was used to estimate the biological processes in which the selected genes were involved.

### Simplified regulation analysis based on sampled fluxes and absolute proteomics

In terms of the fluxes used in protein regulation coefficient computation, Yeast9 were constrained by measured exchange fluxes, total protein concentration, and growth rate. Then, simulated a minimization in uptaking nitrogen source, whereby the minimized value is fixed. Lastly, targeting maximize ATP maintenance, the flux distribution was sampled 1000 times ^57^.

### GEMs reconstruction for single/double knockout *S. cerevisiae* strains in large-scale

The transcriptomic and growth data were originally from previous research ^60,61,82^ collected in the study ^62^. To integrate transcriptomics into Yeast9, the method described in the previous study ^62^ was applied to modify the upper and lower bound of each reaction by gene expression value. From Equation (1), the AND relationship in GPR means the reactions needs all subunits to take place, so the minimum gene expression value of all subunits is used. For Equation (2), the OR relationship describes that only one isoenzyme is able to pull the reaction, so the maximum function is selected.

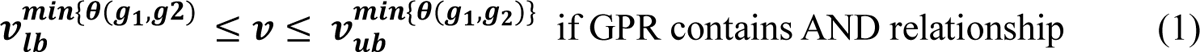

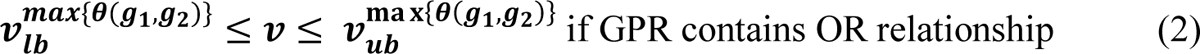

Where θ denotes the gene expression level.

### Gene function prediction using 6 machine learning algorithms

The deleted genes were classified according to the PANTHER classification system ^83^ in a previous study ^62^. In this dataset, the gene number in those functional categories ranges from 29 to 149. Various machine learning models were used to predict the functional categories, to which those metabolic genes in Yeast9 belong. The corresponding predicted fluxes and transcript profiles were randomly divided into training (80%) and testing (20%) subsets. Before training the machine learning models, the data was processed by kPCA, where the components is 500 to extract features ahead of schedule. Using the toolbox of sklearn, random forest, support vector machine, K-nearest neighbour, multilayer perceptron, Naïve Bayes, and logistic regression models were trained using training datasets (Supplementary File 6). Cross-validation over different sets of training/test sets was conducted to evaluate the prediction performances of different machine learning models.

## Availability of data and materials

The Yeast9 is accessible at https://GitHub.com/SysBioChalmers/yeast-GEM and scripts for various omics integrative analysis is available at https://GitHub.com/hongzhonglu/yeast_GEM_multi_omics_analysis. The large file used to construct knockout models is in https://figshare.com/articles/dataset/large_data_used_in_https_github_com_hongzhonglu_yeast_GEM_multi_omics_analysis_/22774076

## Author contributions

HZL and EJK designed the research. CYZ performed the research. LY, CYZ and EJK analysed the data. HZL and EJK supervised the work. BJS, FRL, CWQE, WTS, SNM, UFL, LMB, HGM, MA, ATR, LXZ, JN participated in the curation of the metabolic network. All authors interpreted the results, discussed, drafted and approved the final manuscript.

## Acknowledgements

This work is supported by grant 2022YFA0913000 from the National Key R&D Program of China, Shanghai Pujiang Program, and grant 22208211 and 22378263 from the National Natural Science Foundation of China (NSFC). This work is also supported by the Novo Nordisk Foundation (grant no. NNF20CC0035580); the Knut and Alice Wallenberg Foundation, and the European Union’s Horizon 2020 research and innovation program (grant agreement 720824); National Key Research and Development Program of China (2020YFA0907800); the 111 Project (B18022); Dutch Research Council (Nederlandse Organisatie voor Wetenschappelijk Onderzoek (NWO)) for the UNLOCK initiative (NWO: 184.035.007); DD-DeCaF project; the European Union’s Horizon 2020 research and innovation program (grant agreements 686070 and 720824), CN Yang Scholars Programme; Deutsche Forschungsgemein-schaft (DFG, German Research Foundation) under Germany’s Excellence Strategy—Cluster of Excellence 2186; Centro de Modelamiento Matemático, ACE210010 and FB210005, BASAL funds for Centers of Excellence from ANID-Chile Project ICN2021 044 of the Millennium Scientific Initiative Grant Exploración number 13220002; and CONICYT Becas Chile grant #72180373 (https://www.conicyt.cl/becasconicyt/).. The funding body has no role in the design of the study, analysis and interpretation of the data, preparation of the manuscript, and decision to submit the manuscript for publication.

