## Supplementary figures and images for "Yeast9: A Consensus Yeast Metabolic Model Enables Quantitative Analysis of Cellular Metabolism By Incorporating Big Data"

### Supplementary Figure 1

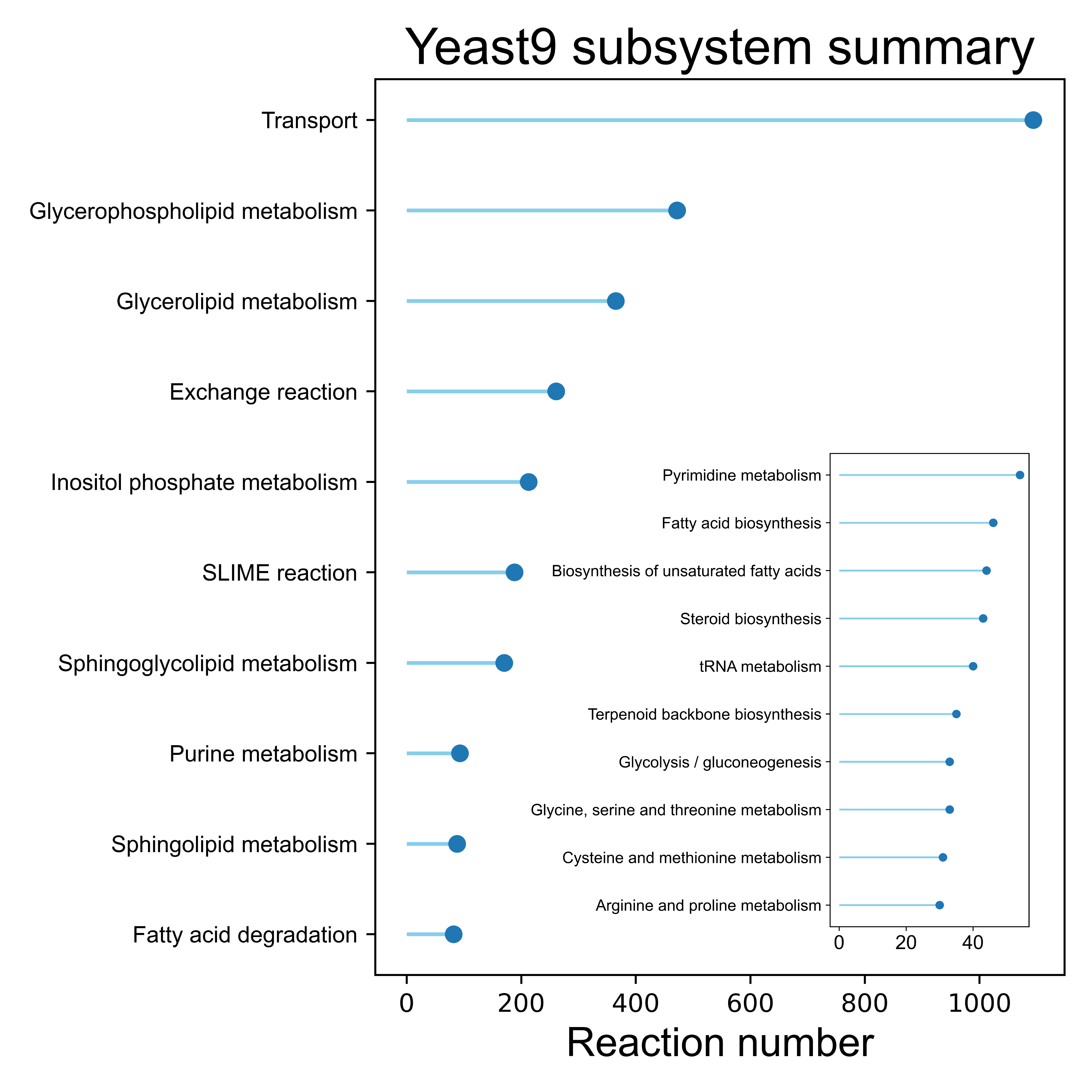

### Supplementary Figure 2

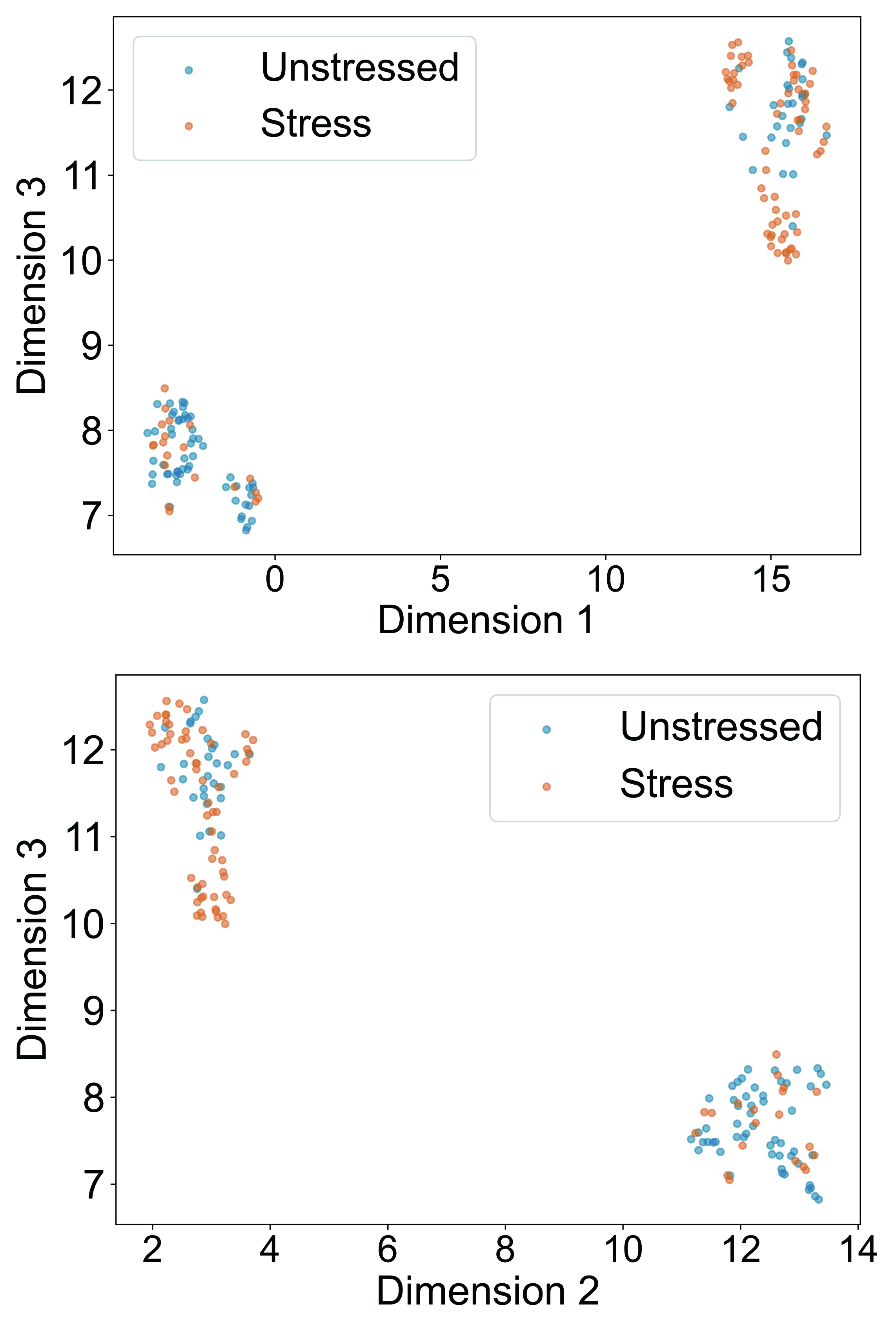

### Supplementary Figure 3

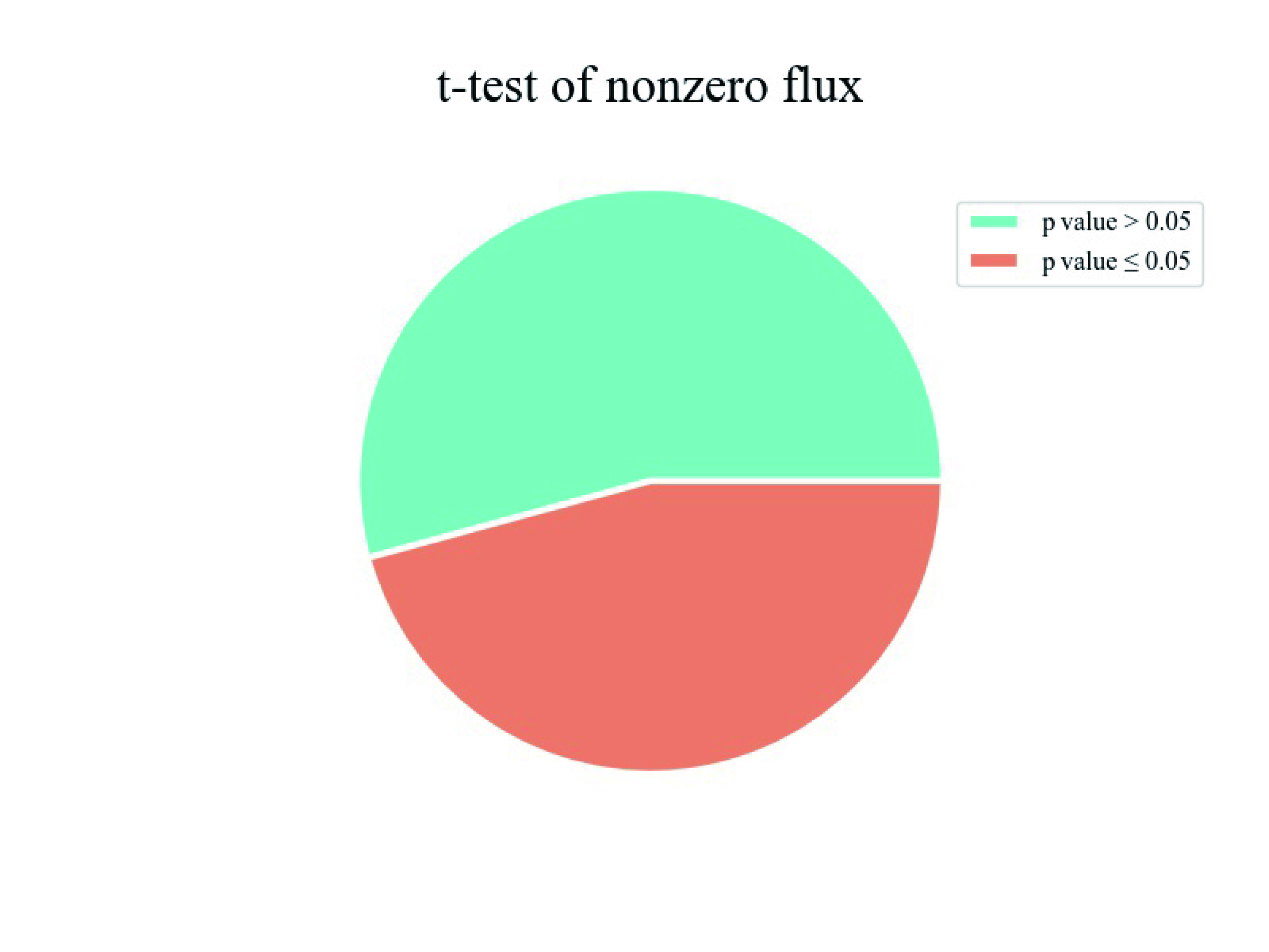
